# Factors influencing tissue cyst yield in a murine model of chronic toxoplasmosis

**DOI:** 10.1101/2022.12.19.521147

**Authors:** Cortni A. Troublefield, Robert D. Murphy, Joy S. Miracle, Ryan W. Donkin, Anthony P. Sinai

**Author notes:** Emergency Department, Good Samaritan Hospital, University of Kentucky Healthcare (Cortni A. Troublefield).

## Abstract

Recent advances into the unique biology of Toxoplasma tissue cysts and the bradyzoites they house necessitates optimization of tissue cyst recovery from infected mouse brains. Here, we present data from 68 tissue cyst purifications of Type II ME49 tissue cysts in CBA/J mice performed over a period of two years. The effects if infecting with both tissue culture tachyzoites as well as *ex vivo* tissue cysts were assessed. Significant mortality was restricted to tachyzoite infections with female mice being more susceptible. Infection with tissue cysts was associated with both lower overall symptomology and mortality exhibiting no sex bias. Cumulatively, host sex did not impact overall tissue cyst yields, although, tachyzoite initiated infections generated significantly higher yields compared to tissue cyst-initiated infections. Notably, serial passage of tissue cysts was accompanied with a decreasing trend for subsequent cyst recovery. The time of tissue cyst harvest, a potential reflection of bradyzoite physiological state, had no significant impact on subsequent cyst yield at the selected time points. In aggregate, the data reveal the considerable heterogeneity associated with tissue cyst yield making the design of adequately powered experiments critical. This is particularly the case for drug studies where overall tissue cyst burden currently serves as the primary and often sole metric of efficacy, as the data presented here demonstrate that cyst recovery between preparations of untreated animals can mirror the reported effects of drug treatment.

## Introduction

*Toxoplasma gondii* is an important opportunistic infection in the context of HIV-AIDS and other immunosuppressive conditions (1, 2). Transmission of the parasite is mediated by two distinct encysted forms, the oocysts, shed at the end of the sexual cycle in the feces of the definitive feline host (3, 4) and the tissue cysts formed with the establishment of the chronic infection in all vertebrate hosts (5). The tropism of tissue cysts to the central nervous system and muscle provides a mechanism of transmission in the act of carnivory (6, 7). Indeed, consumption of raw or undercooked meat contaminated with tissue cysts serves as a primary mechanism for transmission to humans contributing to *Toxoplasma gondii* being a significant agent of food-borne infection (6-8).

Despite their central role in transmission, little is known about the basic biology of the tissue cyst (5). What is known is that the tissue cyst represents a genetically clonal though physiologically heterogenous community of slow growing bradyzoite forms of the parasite (5). Bradyzoites were long viewed to be dormant forms, a position that was challenged by our work demonstrating that tissue cysts derived from infected mouse brains contain bradyzoites exhibiting metabolic activity including the capacity to replicate by endodyogeny (9). This critical finding has triggered a renewed interest in bradyzoite biology as it provides a window into potential drug treatment for a form that remains refractory to currently approved drugs (10).

The development of new treatments targeting encysted bradyzoites will be fundamentally dependent on efficient means to generate tissue cysts in vivo for further functional analyses (10-13). Here we build on our earlier work (9, 14) to identify factors influencing tissue cyst yields from infected brains in a murine model of experimental toxoplasmosis. In seeking to optimize the parameters contributing to tissue cyst recovery from the infected brain, we assess the effect of sex, infection source, the consequence of serial passage of tissue cysts between mice, and the effect of the relative time of harvest on subsequent cyst yield. Additionally, tissue cyst yields were assessed based on the mean recovery per infected animal while also considering per capita yields accounting for animals succumbing to infection prior to harvesting. This latter category proved to represent a significant proportion of animals infected with tissue culture derived tachyzoites.

The data presented herein indicate that tissue cyst yields are impacted by multiple factors necessitating consideration particularly in the interpretation of outcomes of drug studies. Drug efficacy is currently evaluated exclusively based on tissue cyst recovery which has a sensitivity limited to under two orders of magnitude. Thus, even the most effective experimental drugs which reduce recovery by between 80-90% (10-13, 15, 16), fall within the range of cyst recovery in the absence of any drug treatment (5, 9) (and data presented in this study). We therefore need to reassess overall approaches to understanding the chronic infection and focus at the level of encysted bradyzoites, appreciating the heterogeneity of activity within the tissue cyst. Development of these methods, several of which are currently underway in our laboratory will greatly benefit from the optimization of tissue cyst yield, considering the factors investigated here and other factors that remain to be identified.

## Materials and Methods

### Parasite Line

All infections using both tachyzoites and *ex vivo* brain derived tissue cysts utilized a derivative of Type II ME49 parasites with a deletion of the HXGPRT (TGME49_200320) gene (17). This line was generated using CRISPR-CAS9 mediated deletion of the gene (18, 19) followed by selection with 6-Thioxanthine (6TX) (17). Selection of the targeting sgRNA sequences followed published sequences targeting this gene (20). Specifically TgHXGRPT(Exon2, +strand): ATGGTCTCCACCAGTGCTCC; TgHXGPRT (Exon3, -strand): GACAAAATCCTCCTCCCTGG and TgHXGPRT (Exon5, +strand): CTTCTTCGAGCACTTGTCC were introduced into the pSAG1::CAS(-GFP::sgUPRT) plasmid (18, 19) to generate 3 distinct mutagenesis plasmids that were co-transfected into Type II ME49 parasites. Initial enrichment for transfected parasites was achieved using flow cytometry (21) prior to selection with 6-TX at 80mg/ml in αMEM with 7% dialyzed FBS. Following drug selection, cloning and confirmation of the mutation, parasites were maintained by serial passage in HFF cells grown in bicarbonate buffered αMEM supplemented with 7% heat inactivated FBS, 50mM glutamine and 25mM Penicillin/Streptomycin at 37°C in 5% CO_2_. This derivative exhibited identical growth characteristics and *in vitro* switching in response to pH8.2/ambient CO_2_ incubation to wild type ME49 parasites obtained from the AIDS Resource Center (data not shown).

### Mouse infection model

Both male and female CBA/J mice (strain #00065, The Jackson Laboratory, Bar Harbor Maine) were used in these studies. Mice were procured at age 4-6 weeks and habituated in the animal facility for a minimum of 1 week and were provided with standard chow (Teklad 2918 irradiated 18%CP rodent diet, Envigo, Indianapolis, IN) and water *ad libitum* during the course of these studies. Mice were infected with Toxoplasma by intraperitoneal (i.p.) injection typically within 2.5 weeks of receipt from the vendor. Infection with cell culture derived tachyzoites was performed using syringe passaged parasites recovered in PBS and kept on ice. At least two independent counts were used to establish the number of tachyzoites in the seed stock. Innocula containing 100 tachyzoites/ 200μl per mouse (500 tachyzoites/ml) diluted into serum free OptiMEM (Gibco). The inoculating suspension was maintained on ice and 200μl injected i.p. per mouse. Prior to injection mice were mildly anesthetized with 30% isoflurane in propylene glycol using a drop jar method (22).

Infection of animals with brain derived tissue cysts was performed by diluting a reserved volume of brain homogenate obtained in the course of purification of brain derived tissue cysts. Brain homogenate corresponding to 20 cysts diluted to 200μl using OptiMEM were used as the innoculum. Quantification of tissue cyst burdens within brain homogenates was established as previously described below (14).

In this study, in addition to tracking the tissue cyst burden from tachyzoite initiated (T) infection, we monitored the effect of serial passage of tissue cysts on the overall tissue cyst burden. Accordingly, the first cycle of infection initiated from tissue cysts (bradyzoites) generated from a tachyzoite-initiated infection represented the B1 inoculum. Subsequent serial passage of tissue cysts from B1 initiated infections were designated B2, followed by B3 and B4 (Figure 1A,B).

**Figure 1A.**
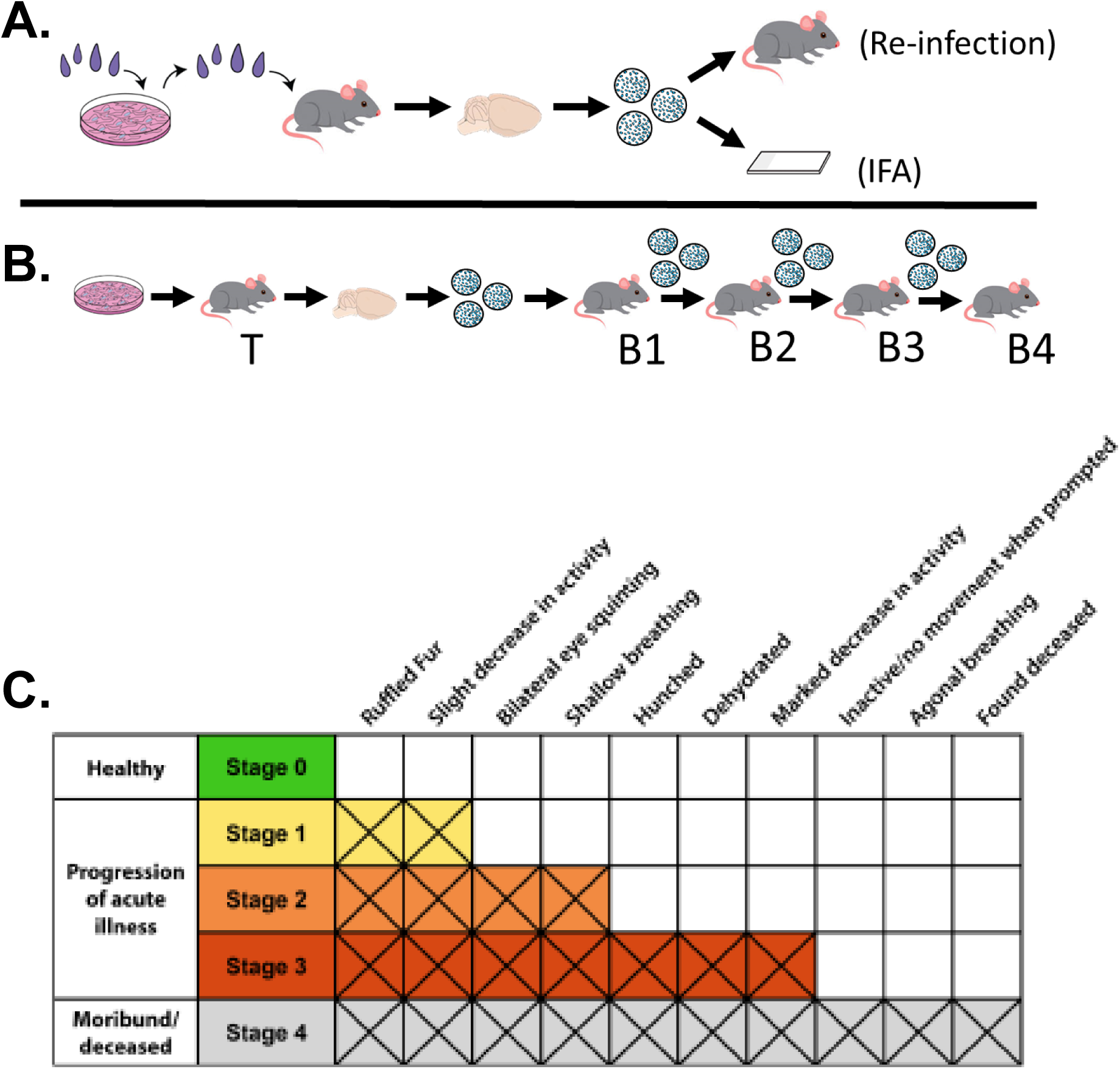
**Schematic of the infection model** initiated with tissue culture derived tachyzoites or tissue cysts harvested from infected mouse brains. Purified tissue cysts were either used for serial passage in CBA/J mice, analyzed by immunofluorescence microscopy (IFA) or used in other applications (purification of RNA for transcriptomic studies). **B**. Sequential passage of tissue cysts derived from an initial tachyzoite (T) infection to establish the 1^st^ (B1), 2^nd^ (B2), 3^rd^ (B3) and 4^th^ (B4) passage. Passages were performed using cysts harvested in Weeks 4,5 or 6 post infection. **C**. Color coded body score index based on symptomology associated with the progression of the chronic infection used to stage disease progression.

Following infection, mice were monitored daily for the development of symptoms with the monitoring increased to at least twice daily once animals became symptomatic. The progression of the infection was monitored using a body score index rubric (Figure 1 C). Notably, this rubric, which reflects the typical progression of symptoms, does not always follow the exact progression, but does provide a framework to standardize the severity of symptoms based on physical observation. As symptoms progressed, supportive treatment was limited to the provision of a gel-based diet (DietGel 76A, Clear H_2_O, Westbrook, Maine) and wet chow on the cage floor. Sub dermal saline injections for animals presenting as dehydrated was provided as additional supportive. No additional interventions were provided to symptomatic animals as the administration of either anti-toxoplasma treatments or anti-inflammatory drugs would alter the progression to the encysted state. All studies involving the use of animals and biohazards were approved by the Institutional Animal Care and Use Committee (IACUC) and Institutional Biosafety (IBC) Committees respectively at the University of Kentucky.

### Tissue Cyst Purification and Quantification

Tissue cyst purifications were performed using a modified Percoll gradient method (9, 14) based on the classic Cornelison protocol (23). Of note, all tissue cyst yields reported in this work were generated from Percoll gradients loaded with the brain homogenates from 2 brains of animals of the same sex. While mixed sex gradients were run, these data are excluded from the current analysis. Additionally, tissue cysts recovered from running single brain gradients using a modified protocol (14) are not reported here as prior work has demonstrated that recovery of cysts from two brain gradients is optimal with significantly lower yields when processing single brain gradients (14). Quantification of cyst burdens was established by the summation of the relative cyst numbers detected in Percoll fractions as previously described (9, 14). A mean yield per mouse was calculated from the total yield of a Percoll gradient, resulting in a single data point from a two-brain preparation.

Quantification of the distribution of tissue cysts within the Percoll gradients was normalized relative to the location of the erythrocytes within the gradients, which was assigned fraction 0. This permits the normalization of distribution of tissue cysts accounting for potential variation in the establishment of the gradients.

### Statistical Analysis

Mouse mortality for each sex following either tachyzoite-initiated or tissue cyst-initiated infection was evaluated using a Kaplan-Meier estimator. Additionally, a graphical representation of disease progression is based on body score index data rubric over the first 28 days of infection was plotted as stacked graph.

The distribution of the data related to tissue cyst yield accumulated over the course of this study was found to be non-parametric. We therefore performed statistical analyses with both the raw data and following a logN-transformation (Ln-transform) of the data. The former data are presented in the form of a scatter plot presenting all data points while the Ln-transformed data are presented as box and whisker plots. All statistical analyses were performed using GraphPad Prism with the specific tests applied indicated within the figure legends.

## Results

### Generation of a Type II ME49ΔHXGPRT line

The generation of a Type II ME49DHXGPRT line for use in this study was undertaken to establish the characteristics of what constitutes a parental (WT) line for subsequent studies targeting specific genes that are ongoing and beyond the scope of the current study. The targeted disruption of the hypoxanthine-xanthine guanine phosphoribosyl transferase gene (TgME49_200320) has been demonstrated to have no impact on growth both in culture and *in vivo* (acute infection) (17, 20, 24). Loss of TgHXGPRT can be positively selected for using the subversive substrate 6-thioxanthine (6-TX) (17, 24). Importantly, restoration of activity in ΔTgHXGPRT lines allows for positive selection using a combination of mycophenolic acid and xanthine (MPA-X) providing a marker for positive selection in subsequent gene targeting/ complementation studies (Fig. 2A.) (17, 24).

**Figure 2A.**
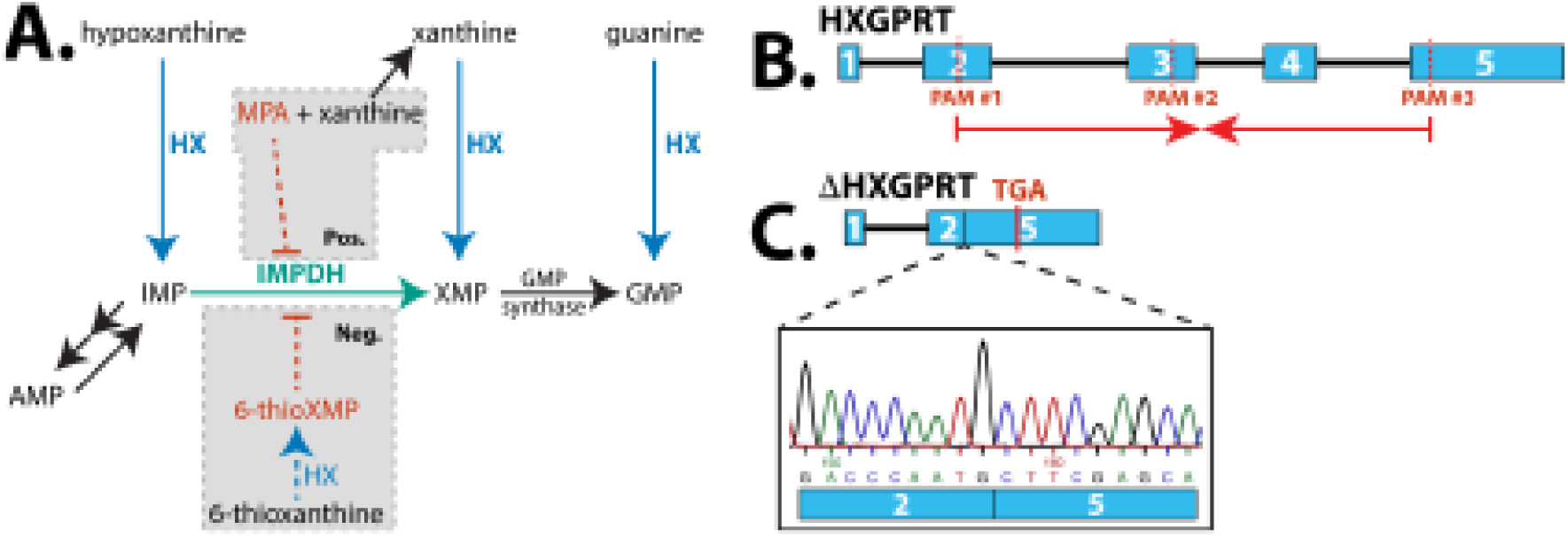
Generation of the ME49ΔTgHXGPRT line. Nucleotide scavenging pathway identifying the basis for the positive and negative selection associated with HXGPRT (HX) KO parasites. Parasites expressing HXGPRT are susceptible to 6-ThioXanthine (6-TX) but resistant to Mycophenolic Acid (MPA) + Xanthine. In contrast ΔHXGPRT parasites are resistant to 6-TX and sensitive to MPA-Xanthine. **B**. Schematic depicting the genomic locus encoding HXGPRT identifying the location of the PAM sites targeted with specific CRISPR-Cas9 constructs. **C**. Schematic of the genomic organization of the ΔHXGPRT locus with the sequence dendogram confirming the fusion of exons 2 and 5 due to the deletion of the intervening introns and exons resulting in the KO phenotype.

To disrupt the HXGPRT locus in the prototypical cyst forming Type II ME49 parasites a shotgun CRISPR-Cas9 strategy (18, 19) was employed as described in the materials and methods. Drug selected parasites were cloned by limiting dilution and the disruption of the gene confirmed by PCR amplification of the locus and sequencing across the mutation lesion (Figure. 2BC). Importantly infectivity and growth characteristics assessed by both replication assays at 24 hours post infection and plaque assays were identical to those of the parental WT ME49 line (data not shown). In addition, the selected clonal TgME49ΔHXGPRT line exhibited identical rates of stage conversion relative to wild type ME49 (data not shown). Finally, this line exhibits no significant difference in the capacity to establish both an acute and chronic infection in mice as detailed extensively below. For these reasons we designate the TgME49ΔHXGPRT parasites, used exclusively in this study as wild type (WT).

### Differential effects of tachyzoite versus tissue cyst infections on mortality

Reports in the literature and our prior experience reveal a considerable heterogeneity in the mortality caused by ME49 tachyzoites during the course of the acute infection (9, 25). These can range from LD_50_ values of 100 to 10^4^ parasites (i.p.), with enhanced virulence selected for by passage in culture. Acute virulence is additionally impacted by mouse genetics which directly or indirectly affect the overall cyst burden (26). In this regard, the CBA/J mouse presents an optimal balance of survival during the acute infection and tissue cyst yields in the brain (9, 26). The recovery of tissue cysts however is inherently highly variable (9, 25) and influenced by a number of factors as discussed below.

In our earlier studies we used female mice exclusively and did not differentiate between tachyzoite or tissue cyst-initiated infections (9). Here, we examined the effect of sex as a variable, as well as the effect of the inoculum on overall survival during the progression of the acute phase and entry into the early chronic phase (days 0-28). Animals were monitored daily and at least twice daily once symptomatic. A body score index, assessing both the physical condition of the animal and its level of activity, was used to assess the progression of the infection (Fig. 1C). Infection with tachyzoites resulted in significant mortality in mice of both sexes, with female mice exhibiting significantly greater levels of mortality compared to males (Fig. 3A). Notably disease progression following tachyzoite infection was rapid with fewer animals surviving despite supportive treatment detailed in the materials and methods (Fig. 3B).

**Figure 3.**
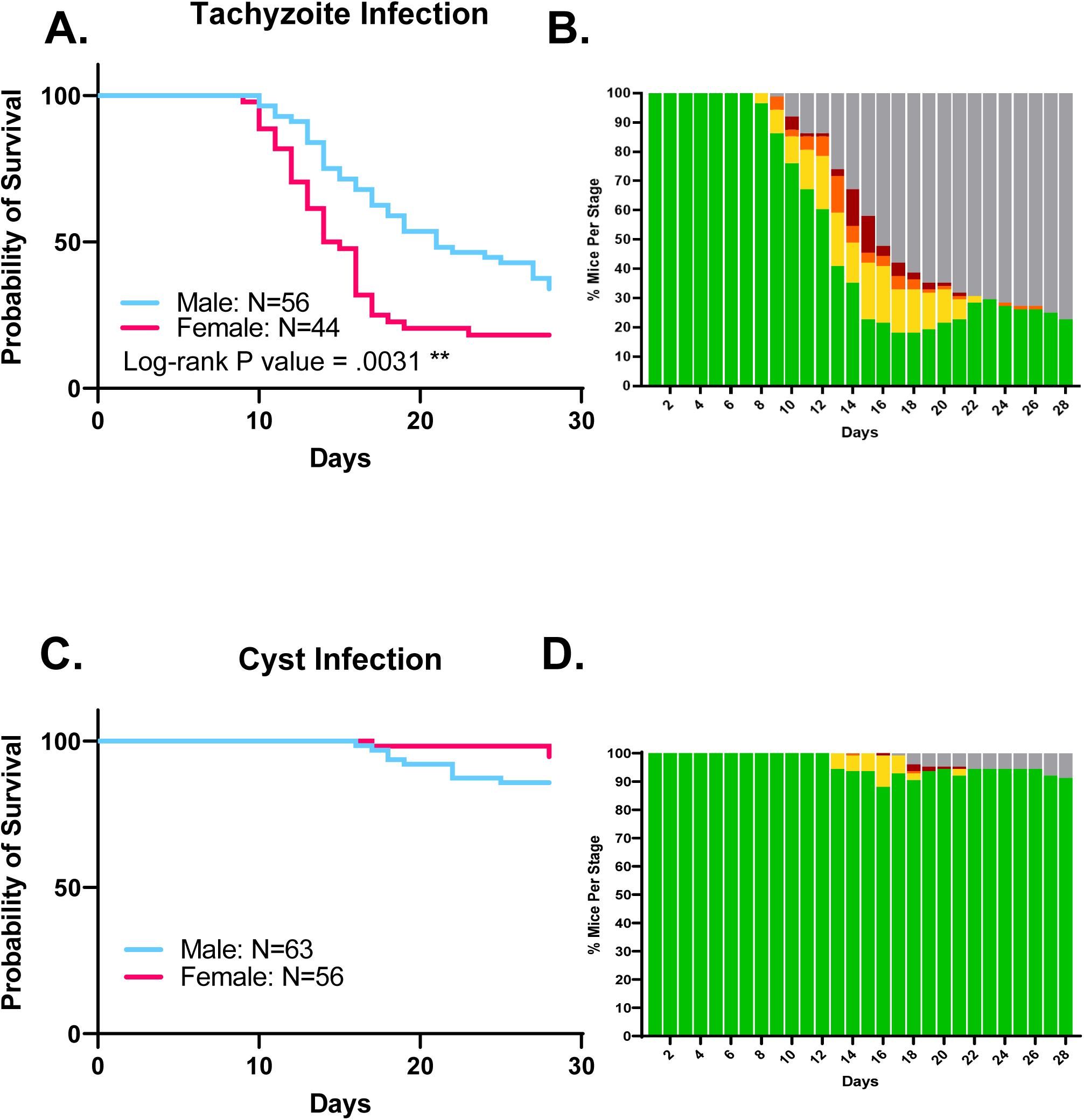
Effect of mouse sex on progression through the acute infection and establishment of the chronic infection. **A**. Infection of male (blue, N= 56) and female (pink, N= 44) mice with 100 tissue culture derived tachyzoites injected i.p. reveal female mice are significantly more susceptible to infection noted by earlier and more extensive mortality compared to infection of male mice. (Log-rank P value = 0.0031 (**\*\***)). **B**. Progression of disease symptomology based on a color coded body score index described in Figure 1C. **C**. Low overall mortality was observed following infection of both male (blue, N=63) or female (pink, N=56) mice with 20 *ex vivo* tissue cysts in brain homogenate. The differences in mortality between male and female mice was not statistically significant. **D**. Low mortality was associated with overall low symptomology scored using a body score index in Figure 1C.

In contrast, relatively low mortality was observed for both sexes with tissue cyst (initiated infections (Figure. 3C). This low incidence of mortality was mirrored by the low overall symptomology as well as temporal delay in the onset of symptoms (Figure 3D). This finding is meaningful in light of the fact that 20 tissue cysts likely possess in excess of 10^5^ parasites (9), a number at least 3 orders of magnitude greater than those causing significant mortality with tachyzoite infections.

### Additional symptomology observed in *T. gondii* infected animals

While the disease progression rubric follows the course of symptomatic disease and recovery during the acute phase and into the chronic infection (Fig. 1C), there were additional adverse events that occurred both during the acute (weeks 1-3) and deep into the chronic infection. The most prevalent of these was the development of a head tilt resulting in animals leaning to one side, sometimes to the extent they exhibited difficulty in lifting their head to feed or drink despite the fact that they often appeared as alert and healthy (Fig. 4). The presentation of head tilts appeared spontaneously with mild head tilts resolving or becoming moderate and severe. Affected animals tended to move in a circular pattern in the direction of the head tilt (clockwise if tilted to the right and counterclockwise when tilted to the left) and varying degrees of a loss of proprioception (Supplemental Data 1). Supportive care was provided, however in most cases animals exhibiting more severe head tilts needed to be euthanized due to their inability to access food or water. During the studies reported here, head tilts (both resolving and progressing) were observed in roughly 20% of animals suggesting they were being triggered by the infection. Among these close to 12% presented with mild or moderate head tilts (13.75% male and 6.25% female) that did not necessitate euthanasia, and 8% (8.75% male and 6.25% female) of animals, with severe head tilts necessitating euthanasia. Notably, there was no observable pattern among the cohorts of infected mice correlating with the development of a head tilt that would be predictive in any way.

**Figure 4.**
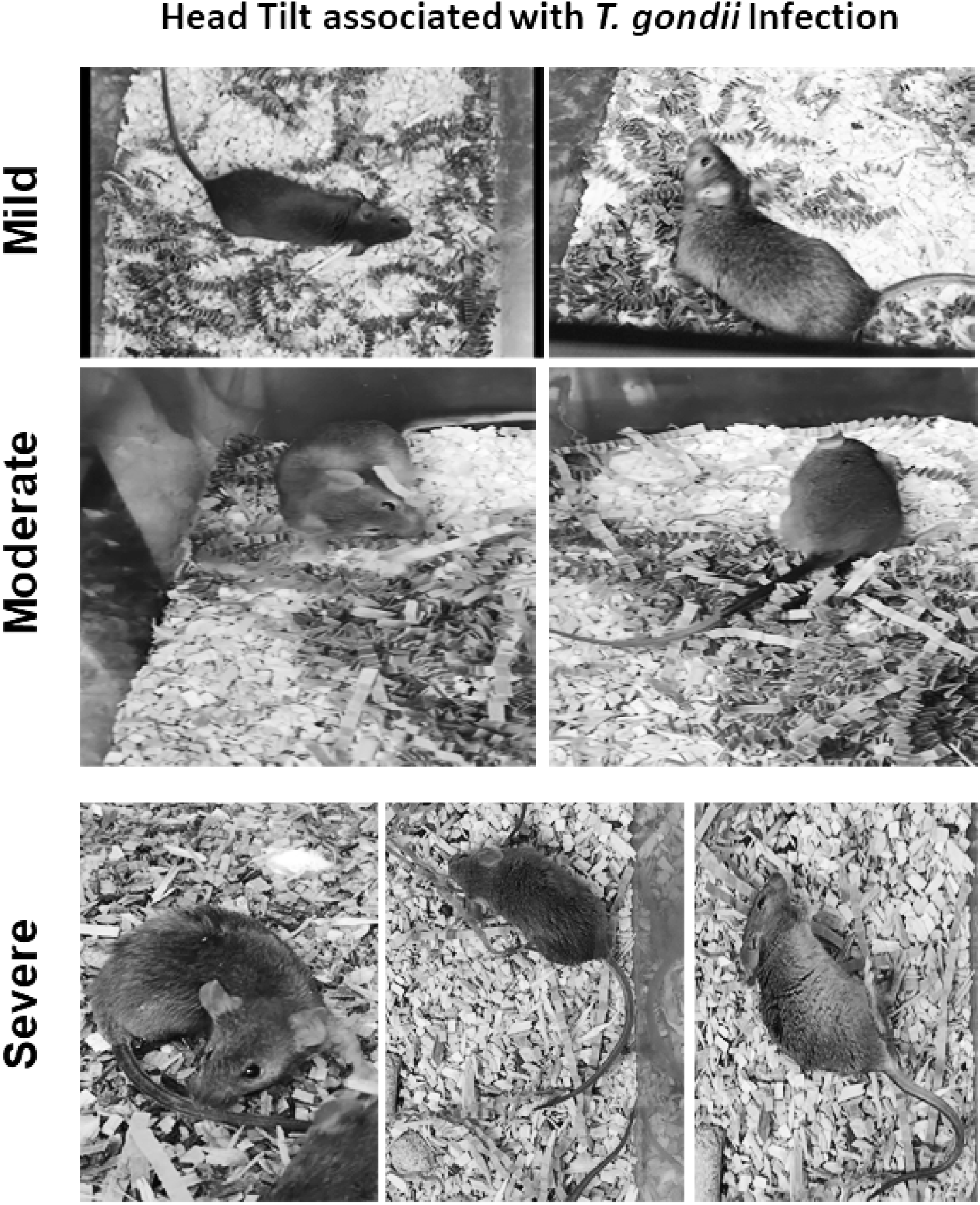
Spectrum of head-tilt presentations in *Toxoplasma* infected mice. A mild head tilt which either spontaneously resolves or progresses to moderate head tilt to a severe head tilt phenotype affecting motility, proprioception and balance. Video recording of motility by head tilt presenting animals is provided in the Supplemental Data.

Consistent with emerging reports (27), infected mice presented with observed seizures both during the acute and chronic phases of infection. In several instances observed seizure were severe enough to warrant euthanasia. It is possible that otherwise healthy looking infected animals that were found dead during the routine daily inspections had succumbed to a seizure. Finally, we observed multiple instances of penile prolapse in male animals which in several instances did not resolve resulting in difficulty with urination and infections necessitating premature euthanasia. With recent reports suggesting colonization of the rodent male genital tract in toxoplasma infections (28, 29), it is conceivable that these symptoms may be connected indirectly with infection.

### Effect of mouse sex and infection type on tissue cyst recovery

The effect of a mouse’s sex on the acute infection, its immune responses to that infection, and other aspects of the chronic infection have been investigated (30-35). Less has been done to establish if the sex of the host animal impacts the recovery of tissue cysts for infections initiated with either tachyzoites or tissue cysts. While sex based differences have been described (30-35), the overall recovery of tissue cysts from infected from male vs female mice infected with either tachyzoites and tissue cysts showed no difference (Fig. 5A,A’). We therefore examined how the inoculum type affected the overall recovery of tissue cysts at multiple time points of harvest. The results clearly indicate that tachyzoite initiated infections consistently generate higher tissue cyst burdens than infections initiated with tissue cysts (Fig. 5B,B’). Of note the lower number of tachyzoite-initiated cyst purifications meeting the criteria for inclusion (2 animals of the same sex-see materials and methods) was due to the high overall mortality associated with tachyzoite infections that precluded an accurate quantification of tissue cysts.

**Figure 5.**
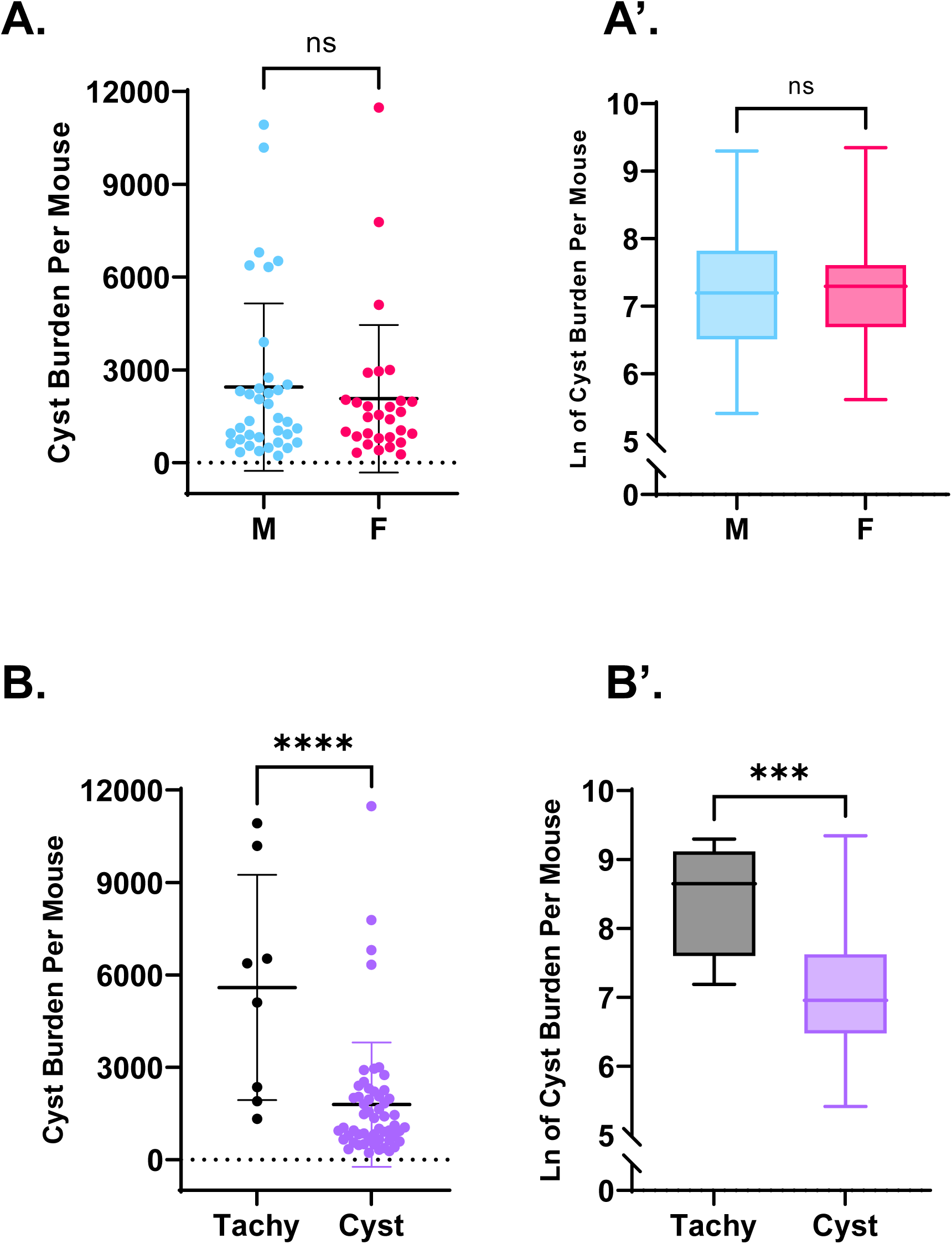
**Mouse sex does not impact overall tissue cyst recovery** based on yields from both male (38 preps) and female (29 preps) mice infected by either tachyzoites or tissue cysts. **A’**. In light of the non parametric distribution of the data statistical analysis was performed on logN-transformed data. These analyses confirm the absence of any statistically relevant differences in tissue cyst recovery as a result of mouse sex. **B**. Tissue cyst recovery as a function of infection with tissue culture derived tachyzoites (Tachy) compared to tissue cysts (Cyst) exposes a significant difference in cyst recovery (Unpaired t test, 2 tailed: P value < 0.0001, ****). **B’**. Analysis of logN transformed data confirms the significant differences in cyst yields between tachyzoite and cyst initiated infections (Unpaired t test, 2 tailed: P value= 0.0001, ***).

### Effect of serial tissue cyst passage on subsequent cyst yields

In light of the lower mortality associated with tissue cyst initiated infections we examined whether serial passage of tissue cysts represented a potential means to mitigate tachyzoite associated mortality (Fig. 6A,A’). The segregation of data based on bradyzoite passage number (B1,B2, B3,B4) confirmed not only the lower cyst recoveries compared to tachyzoite infections but also a trend of diminishing cyst yields. (Fig. 6A,A’). In light of the trend of lower yields with passage, a limited analysis was performed to assess the consequence of infection with cysts following 3 serial passages though mice (B3). The resulting yield for B4 cysts in the low 100’s, close to the level of detection precluded further serial passages (Fig. 6A,A’,B). Together these results suggest that serial passage within CBA/J mice, while exhibiting low overall mortality needs to be balanced with the diminishing yields as quantified on a per mouse infected basis (Fig. 6B).

**Figure 6.**
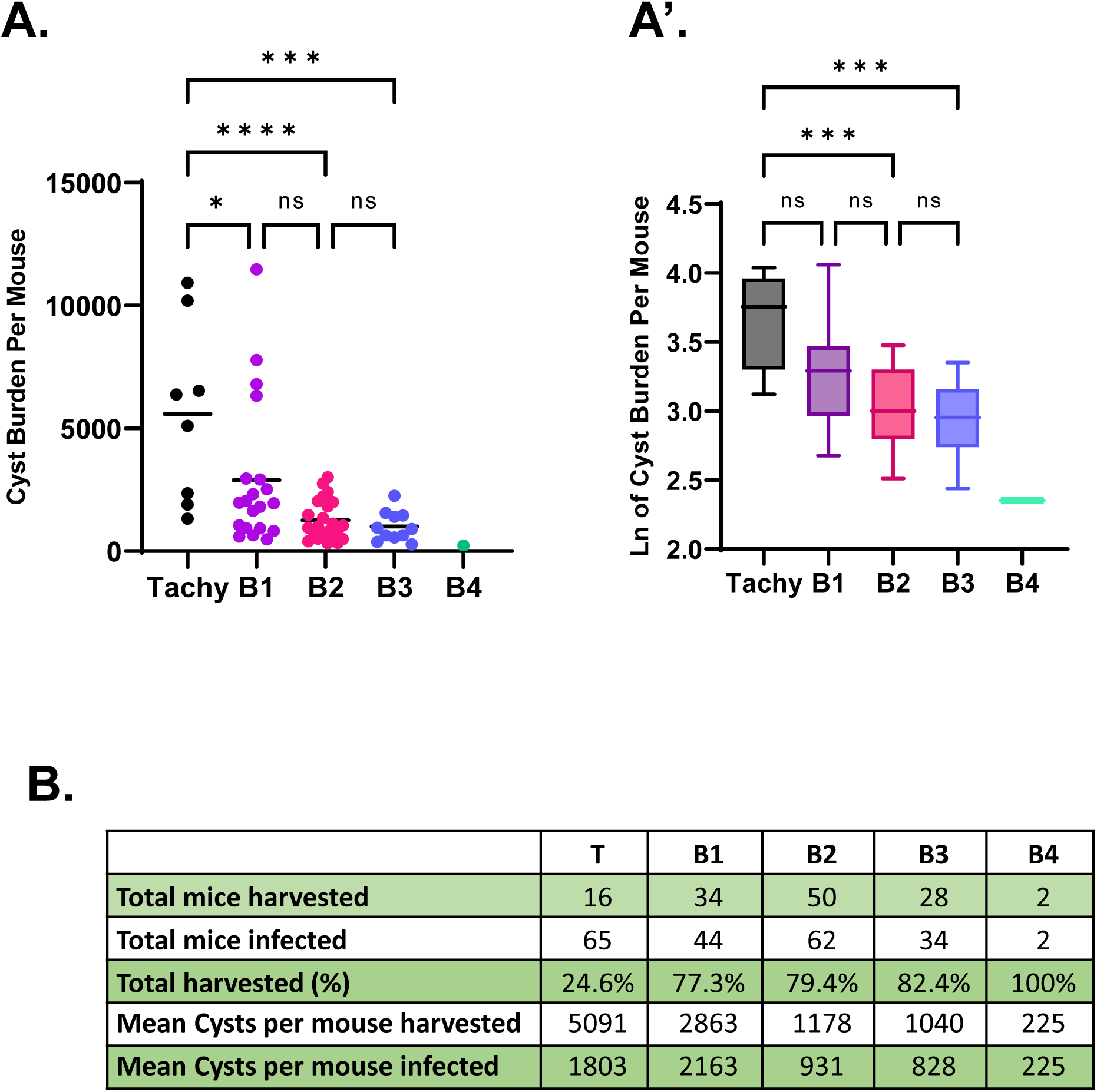
Tissue cyst recovery following serial passage of ex vivo tissue cysts reveals a pattern of diminishing returns. **A**. Segregation of cyst yield data from all tissue cyst initiated infections based in passage number confirms the differences in cyst yield are modest between tachyzoite (T) and the first tissue cyst cycle (B1) (One way ANOVA Tukey’s multiple comparisons test P value= 0.035, *). Significant differences are found between tachyzoite (T) and the second (B2) (One way ANOVA Tukey’s multiple comparisons test P value < 0.0001, ****) and third (B3) (P value = 0.0003, ***) cycles. The single data point for B4 precludes statistical analysis. There was not statistical difference (ns) among the bradyzoite initiated infections (B1, B2, B3, B4). **A’** LogN transformed data reveals no statistical difference between tachyzoite-initiated and B1 cyst initiated infections on subsequent yield. Significant differences in cyst yield were observed when comparing tachyzoite initiated infections with second (One way ANOVA with Tukey’s multiple comparison test P value = 0.0002, ***) and third cycle (B3) cyst infections (P value = 0.0002, ***). Additionally, statistically significant differences were noted between B1 and B3 initiated infections (P value= 0.036, *). **B**. Table indicating the actual recovery of tissue cysts as a function of total animals infected based on infection (Tachyzoite (T) vs cyst (B1, B2, B3, B4). Each prep included uses 2 mice of the same sex. Specific details regarding inclusion criteria are described in the text.

### Effect of time of harvest on subsequent tissue cyst burden

Contrary to long standing dogma, the chronic infection is considerably more dynamic than previously thought (5, 9). Our earlier work (5, 9) suggested that the physical properties of tissue cysts varied, defined by the distribution of their density in Percoll gradients at their times of harvest and is supported in the current data (Fig. 7). These shifts may reflect cyclical changes in the physical properties of the tissue cysts associated with patterns of replication hinted at by the dynamic nature of TgIMC3 (TgME49_244030) labeling, a marker for both active/recent replication (9, 36) and the time since a specific bradyzoite last replicated (9). Such episodic events can result in significant changes on the overall packing density, established by quantifying the actual number of bradyzoites within tissue cysts thereby impacting their density in the gradients (9).

**Figure 7.**
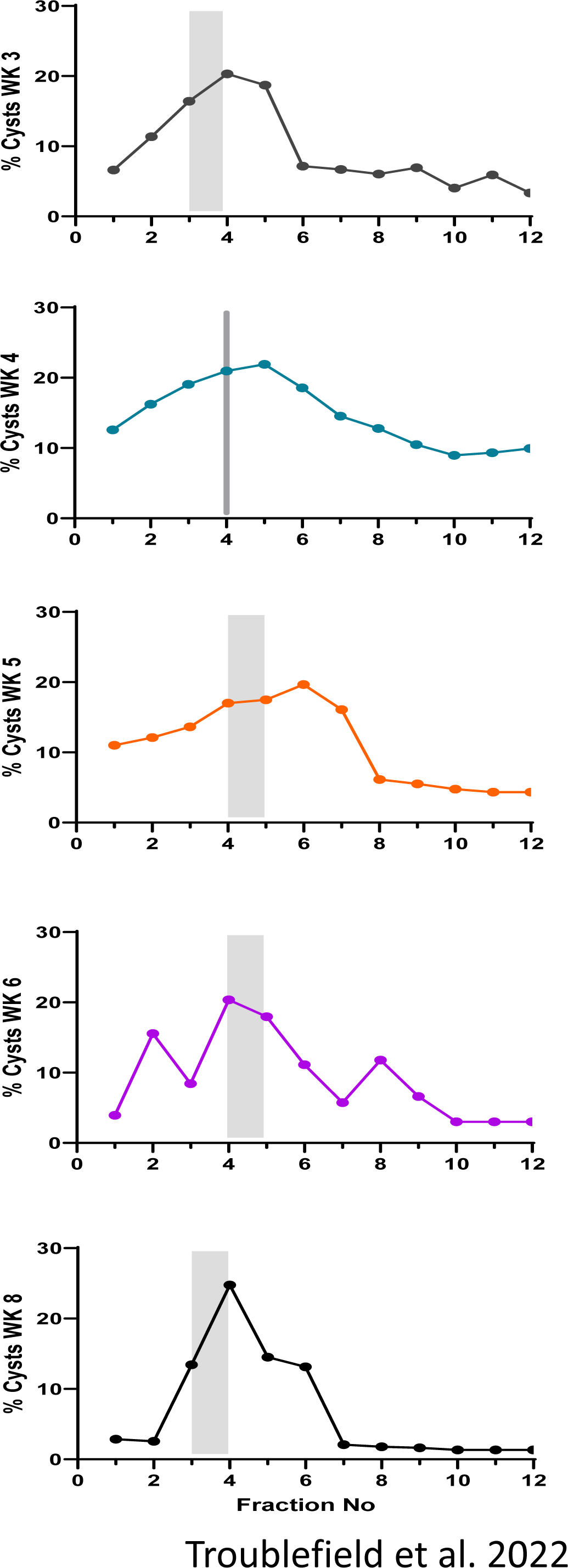
**The physical characteristics of tissue cysts as a function of the time of harvest** post infection reveal a dynamic distribution in Percoll gradients. The mean distribution for the proportion of tissue cysts detected in individual fractions were plotted for each time point (Week 3 n=10, Week 4 n= 23, Week 5 n=13, Week 6 m=13, Week n=9, Week 8 n= 4). The gray shading in the graphs defines the fraction(s) within which the median cyst distributions were recorded. Gradient fractions at the interface of the 90% and 40% Percoll steps representing the fraction concentrated in mouse RBC served as Fraction 0. Tissue cyst distribution in the roughly 1ml fractions (collected a from the 40% step through the brain homogenate overlaid on the 20% Percoll step were plotted as percentage of the total recovered cysts.

TgIMC3 intensity is as a surrogate for the birth-dating of individual bradyzoites that can serve as an indicator of active growth or relative dormancy (9). As such, the proportion of “younger” and more active bradyzoites relative to their inactive peers within the same tissue cyst can potentially impact their capacity to establish a subsequent infections and thereby the overall cyst yield. We therefore reexamined the data for both tachyzoite and bradyzoite-initiated infections resolving whether the time tissue cysts were harvested affected the overall tissue cyst yield in the subsequent infection cycle. What is clear is that while bradyzoite initiated infections as a whole typically yield lower cyst recovery compared to tachyzoite (Fig. 5B,B’), the time of harvest did not significantly impact the subsequent recovery of tissue cysts in subsequent purifications (Fig 8A,A’). Interestingly, infection with tissue cysts recovered at Week 4, a time point earlier in the chronic phase where active replication is projected to be most pronounced (9), produced a large range of subsequent cyst yields. This broad range however was not found to be statistically significant relative to Week 5 initiated cysts, when the lowest metabolic activity is expected (9) and Week 6 cysts where activity is projected to be on an upward trajectory in the cycle (9). Interestingly, tissue cyst yields following infections with B2 and B3 tissue cysts at all “ages” had no impact on subsequent yield but trended lower than recovery following infection with B1 cysts (Fig. 6B).

**Figure 8.**
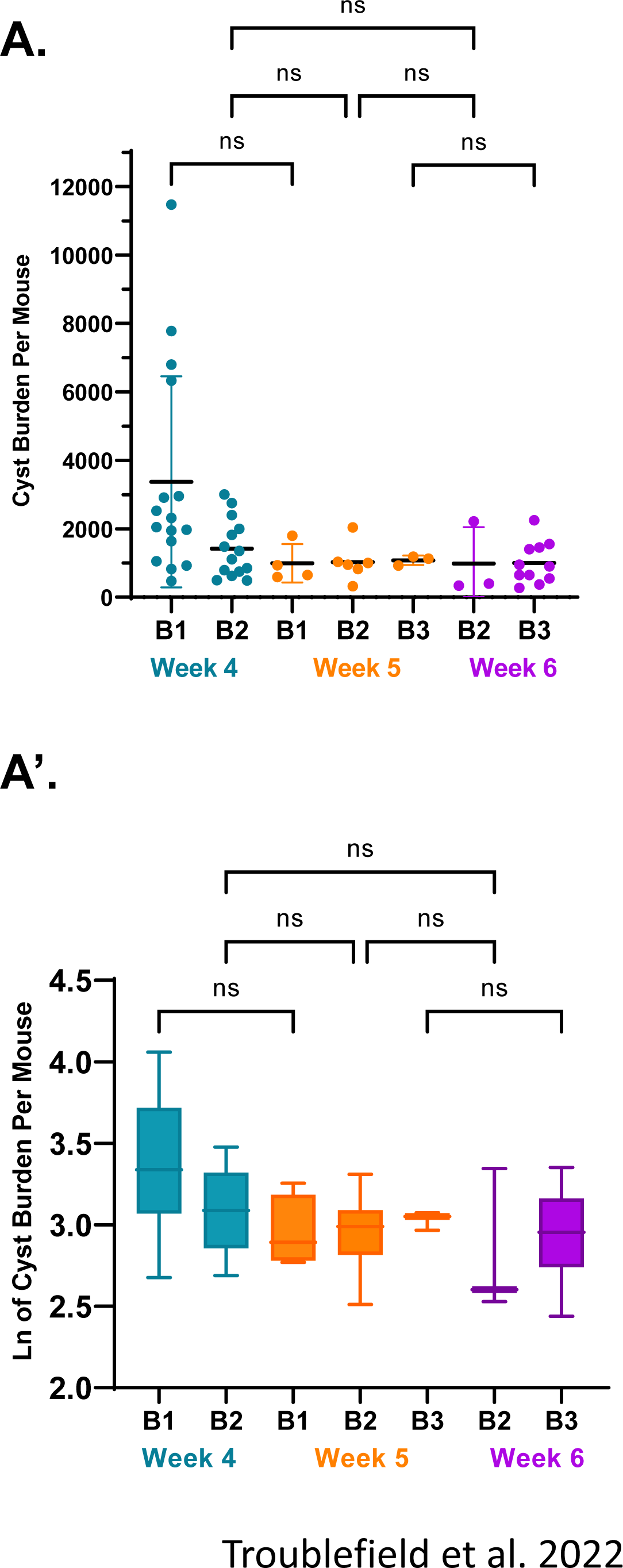
**A. Infection with tissue cysts harvested at Weeks 4, 5 and 6** along with the specific serial passage number (B1, B2, B3, B4) exhibit no differences in sub-subsequent cyst yields despite marked differences in their distributions within Percoll gradients. **A’** LogN transformed data confirms the lack of differences in subsequent cyst recovery as a function of the time of harvest of the infecting tissue cysts.

## Discussion

Bradyzoite within tissue cysts are arguably the most clinically relevant form in reactivated disease upon immune suppression (10, 37, 38). Despite its central role in the pathogenesis, little is known about the biology of the bradyzoite and the community of bradyzoites that represent each individual tissue cyst (5). Notably, while bradyzoites within a tissue cyst are genetically clonal, they represent a physiologically diverse population that varies from cyst to cyst even within the same animal (5, 9). As such, a bradyzoite retains a level of plasticity, permitting it to remain in this state, dedifferentiate into a tachyzoite during recrudescence or enter the sexual cycle in the feline gut (39). It is tempting to speculate that the heterogeneity inherent within the genetically clonal tissue cyst is central to the survival of the organism.

The variable effect of infection in the murine model has been well documented with both parasite effectors and host factors engaged in a complex interplay defining susceptibility and resistance (26, 32, 38, 40-42). Beyond the inherent variability observed with the cyst burden, differences with regard to parasite lines and mouse backgrounds (26), and differences with regard to the sensitivity of approaches used to quantify the cyst burden present vastly different numbers as summarized in a comprehensive review on the topic (25). These differences with the lack of standardization complicate comparisons across studies, which is becoming increasingly important with the renewed focus on the chronic infection. In the context of cyst formation and yield, the pairing of Type II ME49 parasites (and its derivatives, like the ΔHXGPRT line used here) and CBA/J mice is recognized as providing the optimal balance assuring reproducibly high tissue cyst yields (25). We have therefore used this specific pairing but found in the course of the studies presented that establishing the conditions for high tissue cyst yields is considerably more nuanced.

While there is no argument with regard to lethality of the mouse hypervirulent Type I strains (RH, GT1), there is considerable variation in the observed LD_50_ values dependent on the passage history of the parasite and the mouse background used in infection (25). Indeed, *in vitro* passage of ME49 in tissue culture selects for mouse hypervirulence resulting in a 2 order of magnitude or greater reduction in the LD_50_ value (from 10e^4^ to 100 or fewer organisms) (26, 43, 44). This is observed to be the case with the lab adapted ME49, and ME49ΔHXGPRT used here. Tachyzoite initiated infections resulted in considerable mortality upon infection with 100 tachyzoites, with a statistically different susceptibility profile for male vs female animals (Fig. 3A). As such, female mice tended to get sicker earlier and succumb at higher rates relative to males in response to i.p. injection with 100 ME49 tachyzoites during the acute infection (Fig. 3B). Recent work from the Knoll laboratory examining the transcriptional responses within the host during the progression of the chronic infection highlights clear differences between the responses between the sexes (44), despite the overall response profile based on several other criteria being fairly similar. Of note, similar to this study, they used CBA/J mice (albeit of different ages than used here) of both sexes and Type II ME49 parasites maintained in tissue culture. Yet, infection doses of 10^4^ allowed for survival in the acute phase and establishment of the chronic infection (44), highlighting the importance of empirically establishing the specific tachyzoite infection regimen, as sex, the age of the mice as well as additional factors impact the outcome of the acute infection that precedes and is essential to the establishment of the chronic infection being interrogated here.

In a previous study, using exclusively female mice, we established that infection with 20 *ex-vivo* tissue cysts i.p. from infected brain homogenates reproducibly generated robust cyst yields (9). Extending these findings we confirmed that infection with tissue cysts of animals of both sexes is associated with very low mortality or symptomology (Fig. 3C.D.). Despite the absence of overt symptoms during the course of the infection, tissue cysts were recoverable from the brains of infected animals, of both sexes with no sex bias related to yield (Fig. 5A,A’). Differences in overall yield were observed when comparing tachyzoite initiated vs tissue cyst initiate infections (Fig. 5B).

One aspect of the toxoplasma infection, that has not been reported on previously, is the development of a head tilt which in severe cases necessitates euthanasia of the affected mouse (Fig. 4). Examination of available literature indicates that infection-associated head tilts in mice have been noted with bacterial infections of the inner ear (45, 46).

Direct brain/CNS involvement is noted in the case of a murine *Plasmodium berghei* study where it may be linked to intra-cerebral hemorrhages (47). Cerebral hemorrhages are also associated with increased rodent head tilts in an ischemia reperfusion injury model, independent of an infection being present (48, 49). Finally, dysregulation of inflammatory responses resulting in intracerebral inflammatory foci and linked to interferon gamma signaling in a model for autoimmune encephalomyelitis is also associated with increased incidence of a head tilt (50, 51). Further indication for an association between an aberrant inflammatory response is evident from head-tilt mice encoding specific mutation in Nox3, the NADPH oxidase involved in inflammation (52). Finally, the absence of any reports regarding the development of head-tilts in the *T. gondii* murine infection may further indicate differential susceptibility of different mouse strains, whereby the CBA/J animals used here are predisposed to developing this condition.

The recovery of tissue cysts following infection with either tachyzoites of encysted bradyzoites exhibited no sex specific differences (Fig 5A,A’) suggesting that the differences seen following tachyzoite infection were likely due to differences in the host response as opposed to factors intrinsic to the parasite. The more aggressive host response in tachyzoites paradoxically resulted in significantly higher tissue cyst recovery in surviving mice (Fig 3A,B and Fig 6B) compared to the cumulative recovery following cyst infection where lower symptomology and mortality was observed (Fig. 3C,D and Fig. 6). This finding would suggest that maintenance of the infection by serial passage of brain derived *ex vivo* tissue cysts would effectively get around the issue of tachyzoite infection associated mortality which precludes cyst recovery. Unfortunately, our data establish that serial passage of tissue cysts results in a progressive trend toward reduction in overall recovery of cysts (Fig. 6). This indicates the need to re-initiate an infection cycle with tachyzoites, despite high mortality, to sustain cyst numbers for downstream analysis. The trend of decreasing cyst yields is not observed in a ME49 line maintained in the Wilson laboratory by serial passage over multiple years in mice (43). A key difference in that maintenance protocol is the “re-charging” of that line in out bred Swiss Webster animals (43) wherein presumed epigenetic changes are reinstituted to permit the restoration of high cyst yields upon return to infections in inbred mice.

Our prior and ongoing analysis of bradyzoite replication within tissue cysts points to largely asynchronous growth that broadly follows a cyclical pattern at least during the first 8 weeks of infection (9). The inner membrane complex protein TgIMC3 (36) serves as a reporter for both parasite birth-dating as well as their relative age based on the intensity of labeling (9). Thus, early in the chronic phase, at week 3 post infection (TgIMC3 high) parasites are in a higher state of metabolic activity associated with extensive replication, both recent and active (9). At week 5 post infection this activity is universally low (9), recovering to an intermediate level at week 8 (9). We reasoned therefore that infection of mice with more “active’ cysts could result in overall higher cyst yields. This however was not found to be the case (Fig. 8). Rather, for bradyzoite initiated infections, the passage cycle (B1,B2,B3) (Fig. 6, 8) appeared to be a primary determinant of subsequent cyst recovery suggesting that the apparent reprogramming by continuous maintenance in a given mouse background (CBA/J in this case) is a key determinant for tissue cyst initiated infections.

Insights into the physiology of bradyzoites is derived primarily from *in vitro* studies using stress-induced stage conversion (53-56). These studies while important in addressing the machinery of stage conversion do not provide meaningful insights into the actual biology of bradyzoites *in vivo*. What limited studies reveal is that despite being genetically clonal, individual tissue cysts exhibit considerable physiological heterogeneity with regard to the resident bradyzoites (5, 9). Our development of imaging based approaches to quantify bradyzoite numbers within *ex vivo* purified cysts revealed the scope of heterogeneity (5, 9), a feature that is reinforced by ongoing studies with the development of imaging based quantitative physiological readouts, including quantification of, mitochondrial morphology and activity (57), amylopectin and replication potential (Patwardhan and Sinai, unpublished). These tools are revealing insights into bradyzoite physiology that will inform drug discovery and development against this currently intractable life cycle stage.

A fundamental limitation in the evaluation of potential drugs effective in the chronic infection is the fact that efficacy measured by the apparent reduction in cyst burdens following treatment are presented in percentage terms (10, 12, 13, 15, 16, 58). While this level of sensitivity represents the currently acceptable norm, they fail to address the mechanistic basis for drug susceptibility or resistance. Initiating investigation at the level of resident bradyzoites with an appreciation for their heterogeneity will lead to more meaningful insights into the biology of the recalcitrant life cycle stage.

Our findings that variations in cyst yield analogous to what is reported for drug treatment (Fig. 5,6,8) is influenced by nature of the initial inoculum, passage and other yet to be determined factors highlights the inefficiency and potential inaccuracy of using the recovery of tissue cysts as the sole metric of drug effectiveness. These issues can be compounded further when using solely the cyst burden to assign functional attributes to mutations in mutant parasite lines (56, 59, 60). As with studies on drug efficacy, the experimental design has to be sufficiently powered to account for the inherent variability in cyst recovery with wild type or parental parasites. As the field continues to recognize that tissue cysts and the bradyzoites they house are not dormant entities, directed physiologic, metabolomic and transcriptomic studies which will depend on being able to recover high cyst numbers for analysis will become increasingly critical. Our objective in compiling the data in the current study is to highlight factors driving cyst yield all of which are subject to further refinement. We hope this work will encourage further studies in the field aimed at optimizing the recovery of tissue cysts to promote meaningful in-depth studies into this critical and fascinating life cycle stage of the parasite.

## Supporting information

Supplemental Movie 1-connected to Fig. 4

## Acknowledgements

The authors would like to acknowledge the assistance of Dr. Heather Bush and the Biostatistics, Epidemiology and Research Design (BERD) Core at the University of Kentucky for guidance with the statistical analysis of these data. The authors wish to thank the vivarium staff of the Department of Laboratory Animal Research for their support and assistance during the course of these studies. We thank Dr. Aashutosh Tripathi for his feedback regarding this manuscript. This work is supported by grants from the US National Institutes of Health RO1AI145335 and R21 AI150631 awarded to A.P.S..

## Supplemental Data 1

SD1: Video: Presentation of mild, moderate and severe head-tilt in *T. gondii* infected mice:

